# Penalized Logistic Regression Analysis for Genetic Association Studies of Binary Phenotypes

**DOI:** 10.1101/2021.02.12.430986

**Authors:** Ying Yu, Siyuan Chen, Samantha J. Jones, Rawnak Hoque, Olga Vishnyakova, Angela Brooks-Wilson, Brad McNeney

**Affiliations:** Department of Statistics and Actuarial Science, Simon Fraser University, Burnaby, BC, Canada; Department of Biomedical Physiology and Kinesiology, Simon Fraser University, Burnaby, BC, Canada; Canada’s Michael Smith Genome Sciences Centre, Vancouver, BC, Canada

**Keywords:** Rare genetic variants, Penalized logistic regression, log-*F* priors, Monte Carlo EM, Laplace approximation, Data augmentation

## Abstract

**Introduction:** Increasingly, logistic regression methods for genetic association studies of binary phenotypes must be able to accommodate data sparsity, which arises from unbalanced case-control ratios and/or rare genetic variants. Sparseness leads to maximum likelihood estimators (MLEs) of log-OR parameters that are biased away from their null value of zero and tests with inflated type 1 errors. Different penalized-likelihood methods have been developed to mitigate sparse-data bias. We study penalized logistic regression using a class of log-*F* priors indexed by a shrinkage parameter *m* to shrink the biased MLE towards zero.

**Methods:** We propose a two-step approach to the analysis of a genetic association study: first, a set of variants that show evidence of association with the trait is used to estimate *m*; and second, the estimated *m* is used for log-*F* -penalized logistic regression analyses of all variants using data augmentation with standard software. Our estimate of *m* is the maximizer of a marginal likelihood obtained by integrating the latent log-ORs out of the joint distribution of the parameters and observed data. We consider two approximate approaches to maximizing the marginal likelihood: (i) a Monte Carlo EM algorithm (MCEM) and (ii) a Laplace approximation (LA) to each integral, followed by derivative-free optimization of the approximation.

**Results:** We evaluate the statistical properties of our proposed two-step method and compared its performance to other shrinkage methods by a simulation study. Our simulation studies suggest that the proposed log-*F* -penalized approach has lower bias and mean squared error than other methods considered. We also illustrate the approach on data from a study of genetic associations with “super senior” cases and middle aged controls.

**Discussion/Conclusion:** We have proposed a method for single rare variant analysis with binary phenotypes by logistic regression penalized by log-*F* priors. Our method has the advantage of being easily extended to correct for confounding due to population structure and genetic relatedness through a data augmentation approach.

## 1 INTRODUCTION

Standard likelihood-based inference of the association between a binary trait and genetic markers is susceptible to sparse data bias [1] when the case-control ratio is unbalanced and/or the genetic variant is rare. In particular, when data are sparse, hypothesis tests based on asymptotic distributions have inflated type I error [2] and the maximum likelihood estimator of odds-ratios is biased away from zero [3].

The relevance of sparse data bias to genetic association analysis is highlighted by recent work on methods for genome-wide, phenome-wide association studies (PheWAS) of large biobanks. Despite the potential of multivariate methods that jointly analyze phenotypes (e.g., [4]), approaches for PheWAS of biobank-scale data typically reduce the problem to inferences of association between single nucleotide variants (SNVs) and traits, adjusted for population structure and relatedness among subjects *via* a linear mixed model (LMM) [5, 2] or whole genome regression (WGR) [6]. For valid testing of associations between rare binary phenotypes and/or SNVs, SAIGE [2], EPACTS [7] and REGENIE [6] implement an efficient saddle-point approximation (SPA) to the distribution of the score statistic that yields correct p-values.

EPACTS and REGENIE also offer testing and effect estimation based on Firth logistic regression [3, 8], a maximum-penalized likelihood method that uses the Jeffreys prior [9] as the penalty. In addition to valid tests, the Firth logistic regression estimator of the odds-ratio is first-order unbiased. Reliable effect estimates are important for designing replication studies and polygenic risk scores, and for fine-mapping [10, section 2.3].

A variety of alternative penalties have been proposed for logistic regression, offering more or less shrinkage than the Jeffreys prior [11]. Greenland and Mansournia developed penalized logistic regression based on a class of log-*F* priors indexed by a shrinkage parameter *m* [12]. In our context, log-*F* (*m, m*) penalization amounts to assuming that the log-OR parameter *β* for the SNV of interest has a log-*F* (*m, m*) distribution with density

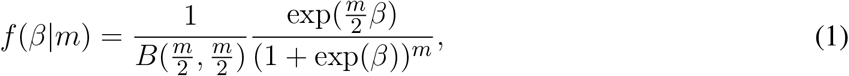

where *B*(·,·) is the beta function (see Figure 1 for plots of log-*F* (1, 1) and log-*F* (10, 10) density curves). In the log-*F* penalization approach, maximizing the posterior density is equivalent to maximizing a penalized likelihood obtained by multiplying the logistic regression likelihood by the log-*F* (*m, m*) prior. The explanatory variables of the logistic regression may include other covariates such as age, sex, genetic principal components (PCs) or the predicted log-odds of being a case from a WGR. In general, we only penalize the SNV of interest but do not penalize other confounding covariates or the intercept, as suggested by Greenland and Mansournia [12].

**Figure 1.**
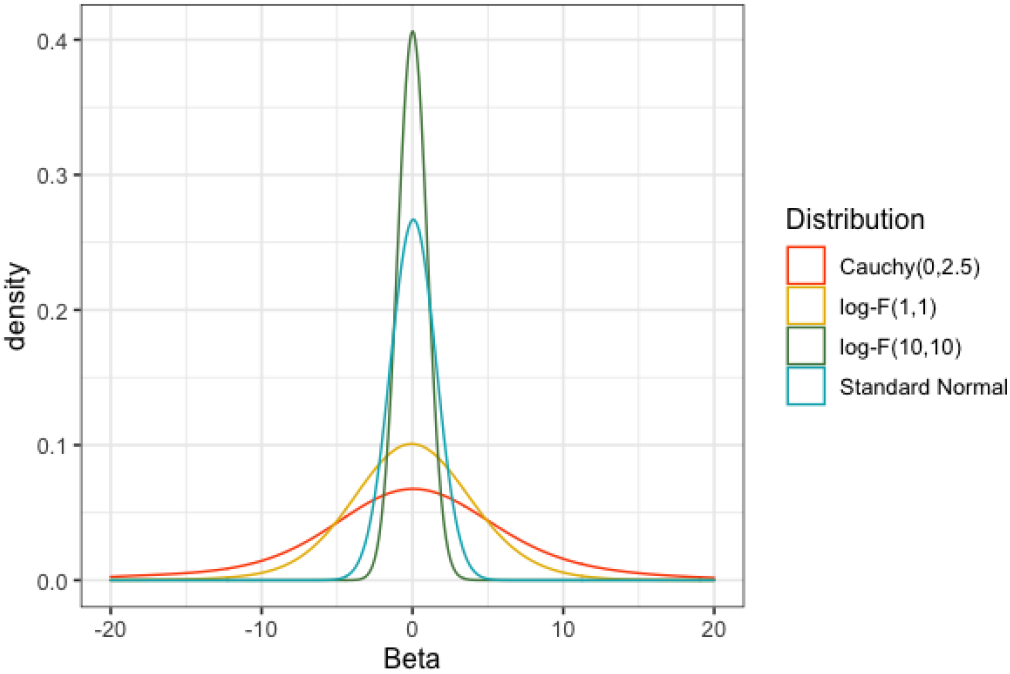
Comparison of log-*F*, standard normal and Cauchy distributions. The log-*F*(*m,m*) density is symmetrically bell-shaped with a single peak at zero, and its variance decreases as increasing *m*. As *m* → ∞, the distribution tends toward a point mass at zero.

Comparisons between log-F-penalized and Firth logistic regression are not straightforward because the log-F approach penalizes selectively, while the Jeffreys prior used in Firth logistic regression is a function of the Fisher information matrix for all coefficients, including the intercept. However, some insight can be gained by comparing approaches for matched pairs data and a binary exposure. For matched pairs, the standard analysis is conditional logistic regression, which eliminates intercept terms from the likelihood. One can show that for a binary exposure Firth-penalized conditional logistic regression is equivalent to imposing a log-*F* (1, 1) prior, which can be implemented by so-called Haldane correction [12]. For Haldane correction we add 1/2 to each of the four cells in the 2 × 2 table of case/control × exposure status and perform a standard analysis of the augmented dataset. More generally, log-*F* (*m, m*) penalized analysis of matched pairs data is equivalent to analysis of the 2 × 2 table with each cell augmented by *m/*2 pseudo-individuals.

Limited simulation studies have shown that, for fixed *m*, log-*F* (*m, m*) penalized methods outperform other approaches for case-control data [11]. Compared to Firth’s method, the log-*F* approach is more flexible, since we can change the amount of shrinkage by changing the value of *m*, and greater shrinkage may reduce MSE [12]. However, there is little guidance on how best to select the value of *m* for a particular phenotype. As a shrinkage parameter, *m* controls the bias-variance trade-off, with the variance of the log-OR estimator decreasing and the bias increasing as *m* increases [12]. We follow the suggestion by Greenland and Mansourinia of using an empirical Bayes method to estimate *m* [12].

Our interest is in fitting single-SNV logistic regressions over a genomic region, or over the entire genome. A motivating example is the Super Seniors study [13] that compared healthy “case” subjects aged 85 and older across Canada who had never been diagnosed with cancer, dementia, diabetes, cardiovascular disease or major lung disease to population-based middle-aged “controls” who were not selected based on health status. The genetic data for this study are described in detail in Section 4. After quality control, data on 2,678,703 autosomal SNVs was available for 427 controls and 617 cases. A preliminary genome-wide scan at a relatively liberal significance threshold of 5 × 10^−5^ found 57 SNVs associated with case-control status.

As in the Super Seniors data, the vast majority of SNVs have little or no effect, and a relatively small set have non-zero effects. The prior used for penalization is the distribution of log-ORs for SNVs with non-zero effects. We therefore propose to select *K* SNVs that show some evidence of having non-zero effects in a preliminary scan, e.g., the *K* = 57 SNVs from the preliminary scan of the Super Seniors data, and use these to estimate *m*. The intent is to learn about the distribution of non-zero log-ORs adaptively from the data [14].

The main goal of this paper is to employ log-*F* penalized logistic regression for analyzing genetic variant associations in a two-step approach. First, we estimate the shrinkage parameter *m* based on a set of variants that show evidence of having non-zero effect in a preliminary scan. Second, we perform penalized logistic regression for each variant in the study using log-*F* (*m, m*) penalization with *m* estimated from step one. For a given *m*, the log-*F* penalized likelihood method can be conveniently implemented by fitting a standard logistic regression to an augmented dataset [12]. In addition to estimates of SNV effects, confidence intervals and likelihood ratio tests follow from the penalized likelihood [8]. Corrections for multiple testing in GWAS/PheWAS applications would involve standard GWAS p-value thresholds, such as 5 × 10^−8^.

## 2 MODELS AND METHODS

We start by reviewing the penalized likelihood for cohort data, followed by the likelihood for case-control data. We then introduce the penalized likelihood and derive a marginal likelihood for the shrinkage parameter *m* based on data from a single SNV. Taking products of marginal likelihoods from *K* SNVs yields a composite likelihood that we maximize to estimate *m*. We conclude by reviewing how log-*F* -penalized logistic regression for the second-stage of the analysis can be implemented by data augmentation.

### 2.1 Likelihood from Cohort Data

Inference of associations between a single-nucleotide variant (SNV) and disease status from cohort data is based on the conditional distribution of the binary response *Y*_*i*_ given the covariate *X*_*i*_ that encodes the SNV. For a sample of *n* independent subjects let ***Y*** = (*Y*_1_, …, *Y*_*n*_) denote the vector of response variables and ***X*** = (*X*_1_, …, *X*_*n*_) denote the vector of genetic covarariates. The likelihood is

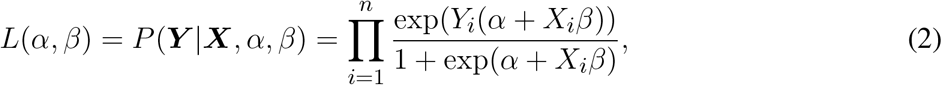

where *α* is an intercept term and *β* is the log-OR of interest.

### 2.2 Likelihood from Case-control Data

The association between a single-nucleotide variant (SNV) and disease status can also be estimated from case-control (i.e. retrospective) data, in which covariates *X*_*i*_ are sampled conditional on disease status *Y*_*i*_ for each individual *i*. Suppose there are *n*_0_ controls indexed *i* = 1, …, *n*_0_ and *n*_1_ cases indexed *i* = *n*_0_ +1, …, *n*, with *n* = *n*_0_ + *n*_1_ denoting the sample size of the study. Qin and Zhang [15] expressed the case-control likelihood in terms of a two-sample semi-parametric model as follows

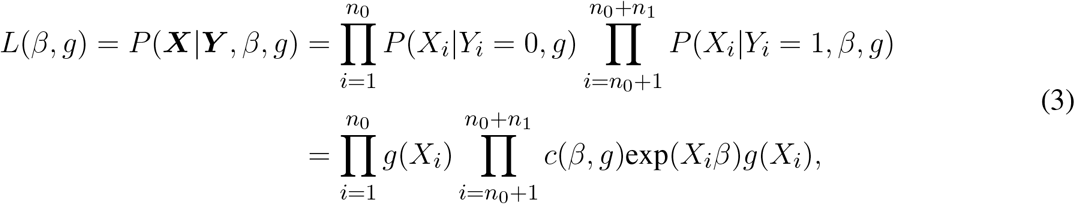

where *c*(*β, g*) is a normalizing constant and *g*(*X*) is the distribution of the covariates in controls, considered to be a nuisance parameter. The potentially infinite-dimensional distribution *g* makes the case-control likelihood *L*(*β, g*) difficult to derive and maximize to find the MLE of *β*. Therefore, we rewrite the case-control likelihood as a profile likelihood [16]:

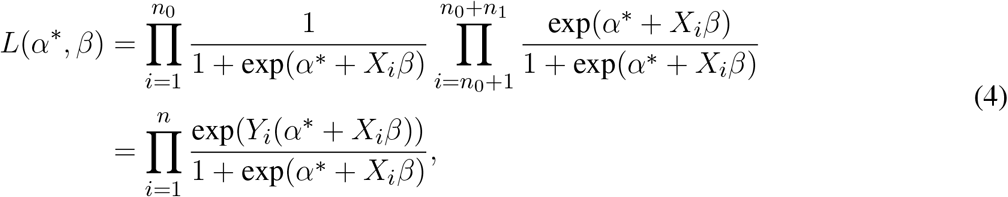

where 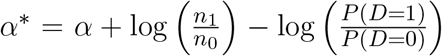, *α* is the intercept term in the logistic regression model for *P* (*Y* = 1|*X*), and *P* (*D* = 1) and *P* (*D* = 0) are the population probabilities of having and not having the disease, respectively [17]. The profile likelihood *L*(*α*\*, *β*) for case-control data is of the same form as the prospective likelihood. The MLE of *β* under the case-control sampling design can be obtained by maximizing *L*(*α*\*, *β*) as if the data were collected in a prospective study [16, 15]. In what follows we write the likelihood as in equation (4) with the understanding that *α*\* = *α* for cohort data.

### 2.3 Penalized and Marginal Likelihoods

The penalized likelihood is obtained by multiplying the likelihood by a log-*F* (*m, m*) distribution (equation (1):

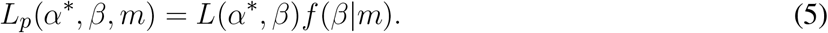

Integrating out the latent log-OR *β* gives a marginal likelihood of *α* and *m*:

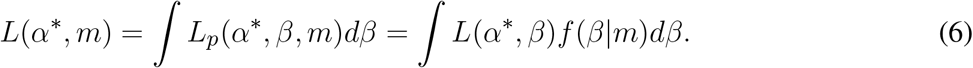

In the above likelihood, the smoothing parameter *m* is the parameter of interest, while the intercept *α*\* is a nuisance parameter.

We expect very little information about *m* in data from a single marker, because this represents a single realization of *β* from the log-*F* (*m*.*m*) prior. In fact, empirical experiments (not shown) suggest a monotone, completely uninformative likelihood roughly 60-70 percent of the time. We therefore consider combining information across markers.

### 2.4 Composite Likelihood for Estimating *m* with *K* markers

Suppose there are *K* SNVs available for estimating *m* (see subsection 2.4.1). For each SNV we specify a one-covariate logistic regression model. Let ***X*** denote a design matrix containing all *K* SNVs, and ***X***_.*k*_, *k* = 1, …, *K*, denote the genotype data on the *k*th SNV. Let 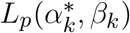 denote the likelihood (4) for the *k*th log-OR parameter *β*_*k*_. Here 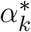 is the intercept term from the *k*th likelihood, considered to be a nuisance parameter.

A composite likelihood [18, 19, 20] for 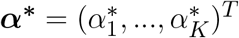 and *m* is the weighted product

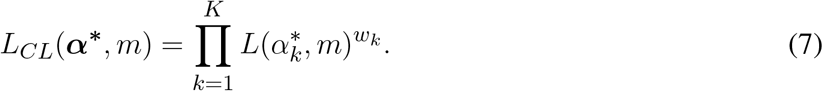

The corresponding composite log-likelihood is

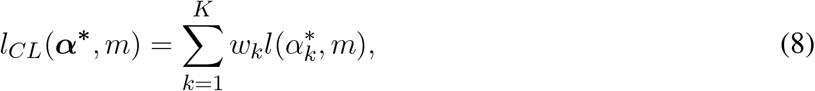

where 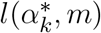 is the marginal log-likelihood contribution from the *k*th variant obtained by integrating *β*_*k*_ out of the joint distribution of observed data and the parameter. Our estimate of *m* is the value that maximizes the composite log-likelihood equation (8). Following the notion that common variants should tend to have weaker effects and rare variants should tend to have stronger effects, we set 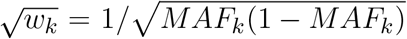 so that *w*_*k*_ is inversely proportional to the MAF of the *k*th SNV [21]. The idea is to up-weight rarer variants of potentially greater effects and down-weight more common SNVs that may have smaller effects.

Maximization is done in two stages

1. For fixed *m*, we maximize *l*_*CL*_(***α***\*, *m*). The form of the composite likelihood when *m* is fixed, as a sum of terms involving only a single parameter, implies that to maximize *l*_*CL*_(***α***\*, *m*) we maximize each 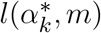 over 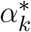. Let 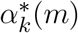 be the value of 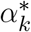 that maximizes 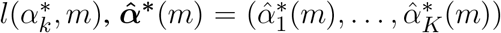, and 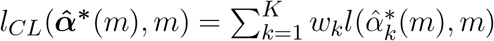.
2. Maximize 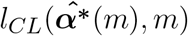 over *m*. To keep computations manageable, we restrict *m* to a grid of values, *m* = 1, 2, …, *M*. One may optionally smooth the resulting 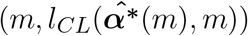 pairs and maximize this smoothed curve to obtain the estimate 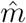.

For a fixed value of *m* and *k*, the estimate 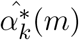 can be obtained by maximizing 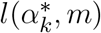 with respect to 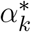. However, it is difficult to evaluate the integral 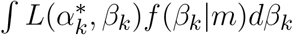 in (6). We discuss two approximate approaches. The first (Section 2.5.1) is a Monte Carlo EM algorithm [22], and the second (Section 2.5.2) is a Laplace approximation to 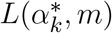 followed by derivative-free optimization of the approximation.

#### 2.4.1 Selecting variants for the composite likelihood

Using variants with no effect in the composite likelihood leads to large estimates of *m*, which correspond to strong shrinkage toward zero. Over-shrinkage biases the log-*F* -penalized estimator towards zero, and reduces power in the second stage of analysis. In the extreme, the use of weakly-associated variants in the first stage can lead to a monotone marginal likelihood in *m* (results not shown). To avoid over-shrinkage we select SNVs with large marginal effects (i.e., small p-values) from a genome-wide scan, similar to the SNV-selection process used by FaST-LMM-Select [23]. For example, we can conduct a preliminary GWAS on all markers, or a thinned set of markers, and choose the SNVs with p-values below a multiple-testing-corrected threshold (refer this as Level 0 of Step 1). We then use the chosen SNVs to estimate *m* (Level 1 of Step 1).

#### 2.4.2 Adjustment for confounding variables and offsets

We conclude this subsection by noting that it is possible to generalize the marginal likelihood approach for estimating *m* to incorporate non-genetic confounding variables, denoted *Z*, and known constants in the linear predictor, or “offset” terms, denoted *b*. As confounders, *Z* will be correlated with the SNV covariates *X*_*k*_, and such correlation may differ across SNVs. We therefore introduce coefficients *γ*_*k*_ for the confounding variables in the logistic regression on the *k*th SNV. Offset terms can be used to include estimated polygenic effects in the logistic regression [6]. Expanding the 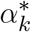 component of the logistic model to 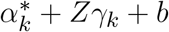, the *k*th likelihood is now

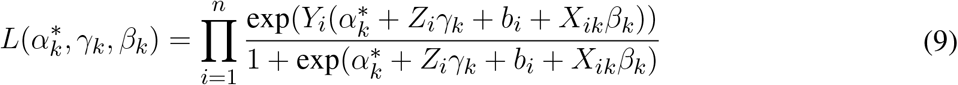

and the composite log-likelihood for estimating *m* is

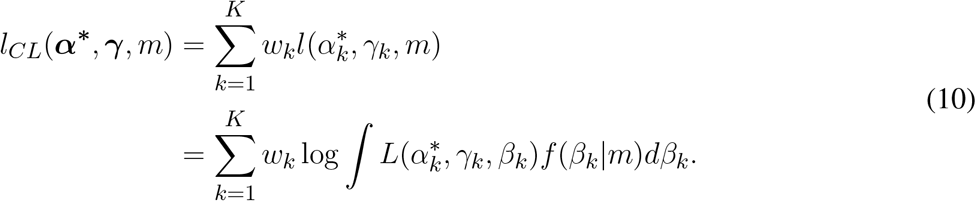

For fixed *m* we maximize *l*_*CL*_(***α***\*, ***γ***, *m*) by maximizing the component marginal likelihoods 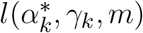 over the nuisance parameters 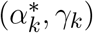. We then maximize the resulting expression over *m* to obtain 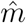. Though the generalization to include confounding variables and offsets is conceptually straightforward, we omit it in what follows to keep the notation as simple as possible.

### 2.5 Maximization Approaches

#### 2.5.1 Monte Carlo EM Algorithm

To maximize 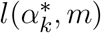, we first consider an EM algorithm, which treats *β*_*k*_ as the unobserved latent variable or missing data. For a fixed value of *m* and *k*, the EM algorithm iterates between taking the conditional expectation of the complete-data log-likelihood given the observed data and the current parameter estimates, and maximizing this conditional expectation. The conditional distribution of *β*_*k*_ given the observed data is a posterior distribution that is proportional to the likelihood *L*(*α*\*, *β*_*k*_) times the prior *f* (*β*_*k*_|*m*). Thus, at the (*p* + 1)^*th*^ iteration, the E-step is to determine

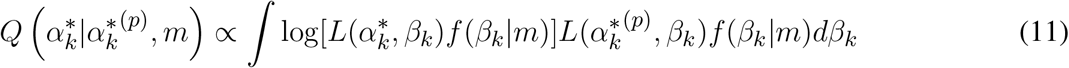

and the M-step is to set

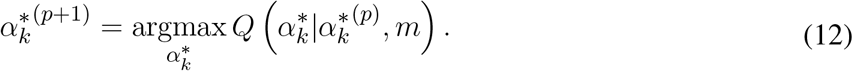

The E-step (11) is complicated by the fact that the integral cannot be solved analytically. We therefore approximate the integral numerically by Monte Carlo (MC); that is, we use a Monte Carlo EM (MCEM) algorithm [24]. The MC integration in the E-step is obtained by sampling from the prior distribution *f* (*β*_*k*_|*m*) [24, 25]. Based on a sample *β*_*k*1_, …, *β*_*kN*_ from the distribution *f* (*β*_*k*_|*m*), the MC approximation to the integral is

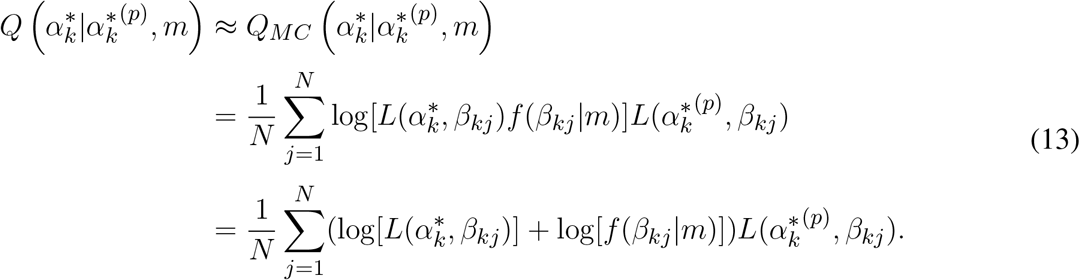

Note that log[*f* (*β*_*kj*_|*m*)] is independent of the parameter 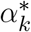, so maximizing (13) in the M-step is equivalent to maximizing

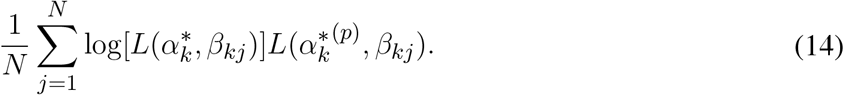

For a discussion of computational approaches to the M-step see the online Supplementary Material.

#### 2.5.2 Maximization of a Laplace Approximation

An alternative to the EM algorithm is to make an analytic approximation, 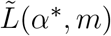, to 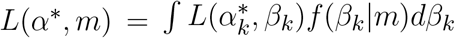 and maximize this approximation. We considered Laplace approximation because it is widely used for approximating marginal likelihoods [26]. The Laplace approximation of an integral is the integral of an unnormalized Gaussian density matched to the integrand on its mode and curvature at the mode. Letting 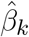 denote the mode of 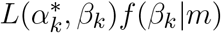 and 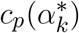 minus its second derivative at 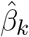, the Laplace approximation to 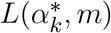 is

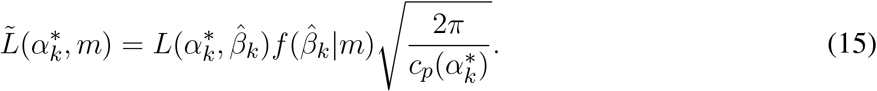

Each 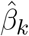 is the root of the derivative equation 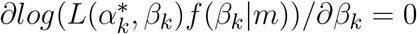; this can be shown to be a global maximum of 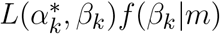. An expression for 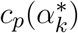 is given in Appendix A of [27]. Figure 2 shows the quality of the LA for one simulated dataset generated under *m* = 4. The approximate marginal likelihood 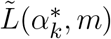 may be maximized over *α*\* using standard derivative-free optimization methods, such as a golden section search or the Nelder-Mead algorithm.

**Figure 2.**
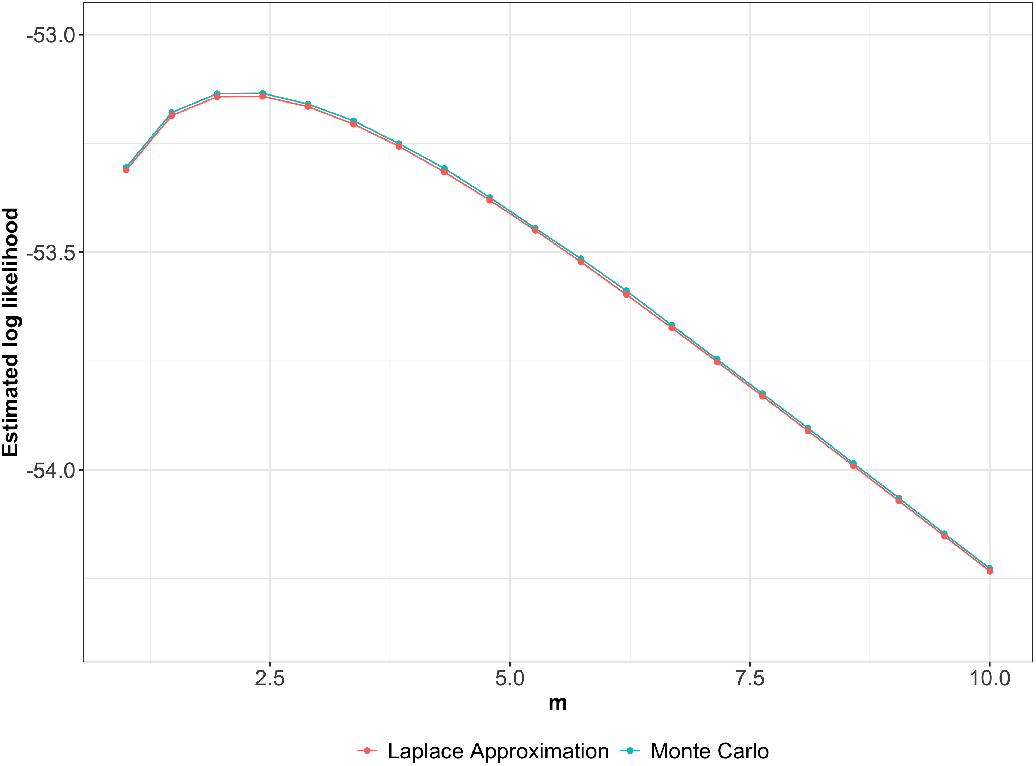
Natural logarithms of estimates of the marginal likelihood 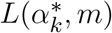 for one simulated dataset generated under *m* = 4. Estimates are obtained by LA and Monte Carlo. Log-likelihood estimates are plotted over the grid *m* = (1, 1.5, …, 10) with 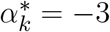.

### 2.6 Implementing log-F Penalization by Data Augmentation

Penalization by a log-*F* (*m, m*) prior can be achieved by standard GLM through data augmentation suggested by Greenland and Mansournia [12]. Here, we provide some details. The logistic likelihood penalized by a log-*F* (*m, m*) prior (equation 5) is:

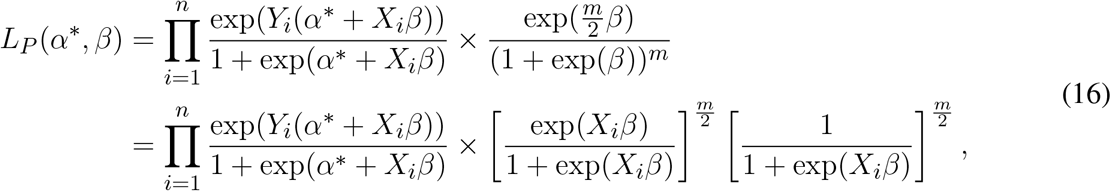

where *X*_*i*_ = 1. Thus, the penalized likelihood *L*_*p*_(*α*\*, *β*) is equivalent to an unpenalized likelihood obtained by adding *m* pseudo-observations to the response with no intercept and covariate one, in which *m/*2 are successes and *m/*2 are failures (even if *m* is an odd number).

In our analyses (see Section 3), we analyze one SNV at a time using the log-*F* penalized logistic regression, adjusting for other confounding variables. The data augmentation approach is illustrated in Figure 3. Let *X* denote the allele count of a SNV and *Z*_*j*_, *j* = 1, …, *p*, denote other confounding variables for adjustment. In the augmented dataset, the response is a two-column matrix with the number of successes and failures as the two columns. The *m* pseudo-observations are split into *m/*2 successes and *m/*2 failures. We only penalize the coefficient associated with the SNV, so we add a single row to the design matrix consisting of all zeros except for a one indicating the SNV covariate. Analyzing the augmented dataset with standard logistic regression yields the penalized MLE and its standard errors, as well as penalized likelihood ratio tests and penalized-likelihood-ratio-based confidence intervals. We conclude by noting that, for fixed *m*, the influence of the *m* pseudo-observations on the fitted logistic regression diminishes as the sample size increases. In other words, for any *m*, the extent of penalization decreases with sample size.

**Figure 3.**
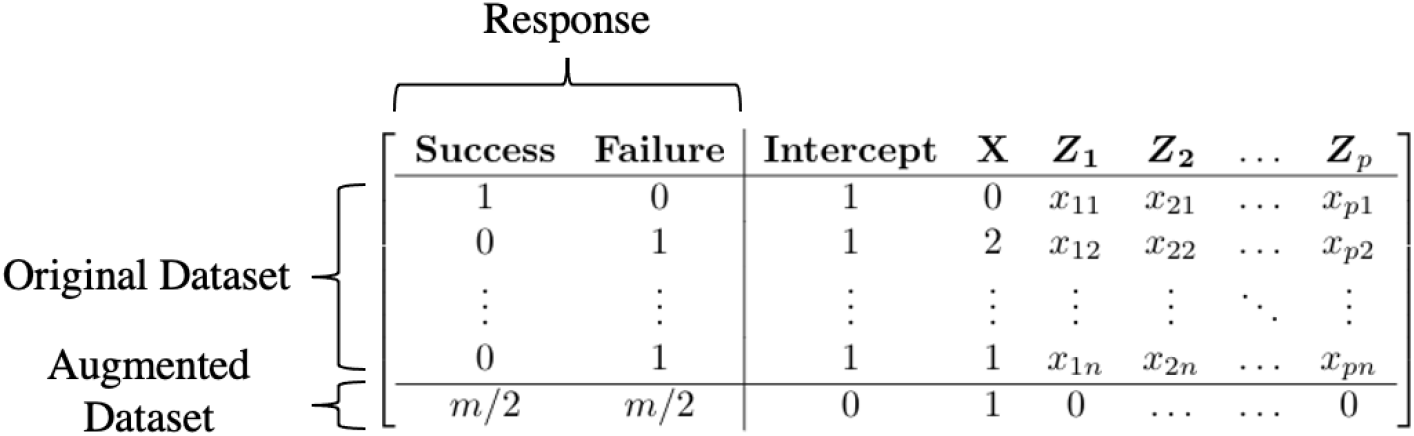
Illustration of data augmentation in the implementation of log-*F*(*m, m*) penalization.

## 3 A SIMULATION STUDY

The empirical performance of the methods introduced in Section 2 was evaluated in a simulation study. The proposed two-step log-*F* -penalized method (LogF) was compared with the standard MLE and the following methods:

- Firth logistic regression (FLR) was first proposed by Firth [3], where the logistic likelihood is penalized by |*I*(*β*)|^1*/*2^ with 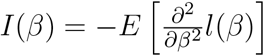 defined as the Fisher information. FLR is implemented in the `R` function `logistf` of the package `logistf` [28].
- Penalization by Cauchy priors (CP) was proposed by [29]. The input predictors are rescaled to have a mean of 0 and a standard deviation of 0.5. All predictors are penalized by a Cauchy prior with center 0 and scale 2.5, whereas the intercept is penalized by a weaker Cauchy prior with center 0 and scale 10. CP is implemented in the `R` function `bayesglm` of the package `arm` [29].

All simulations were performed using `R` (Version 4.1.2) [30] on the Compute Canada cluster Cedar. We restricted *m* to a grid of values between 1 and 10, and we used parallel processing that splits the computation of the composite likelihood for each *m* ∈ [1, 10] over different cores. Each node on the cluster has at least 32 CPU cores and we allocated 10G to each core. For detailed description of its nodes’ characteristics please refer to https://docs.computecanada.ca/wiki/Cedar#Node_characteristics.

We set the sample size to 500, 1000 and 1500, and 100 data sets were generated in each scenario. For each data set, we first estimated *m* based on a set of SNVs which show non-zero effects in a preliminary scan (Step 1), and then implemented the log-*F* penalized likelihood method to test single-variant association for each SNV by the data augmentation approach (Step 2). For the MLE and CP approaches we used Wald tests for SNV effects and Wald confidence intervals for the SNV coefficient. For FLR and the LogF approaches tests we used likelihood-ratio tests (LRTs). For a penalized log likelihood *l*_*P*_ (*α, β*), the likelihood ratio statistic [8] is

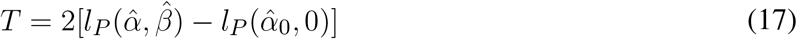

where 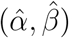 is the maximum of the penalized likelihood function and 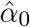 is the maximum of the penalized likelihood when *β* = 0. The p-value is computed from the 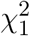 distribution. For penalized logistic method, profile penalized likelihood (PPL) confidence intervals have shown to have better empirical properties than standard Wald-based confidence intervals [8]. A PPL confidence interval can be obtained by inverting the LRT, i.e., by finding all values of 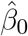 such that 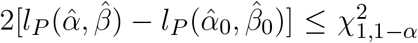 gives a 100(1 − *α*)% confidence interval for *β*.

### 3.1 Data Generation

To keep computations manageable, we simulate preliminary datasets of 50 causal and 950 null SNVs. Null SNVs were used to assess the Type I error performance and the power was estimated using the set of causal SNVs. The data was simulated according to a case-control sampling design, where covariates are simulated based on disease status. For a given SNV, let *X*_*j*_ denote the allele count (i.e. 0, 1 or 2) of SNV_*j*_ and *β*_*j*_ be the corresponding log-OR parameter. Following [15], the conditional density function for *X*_*j*_ in the controls and cases are

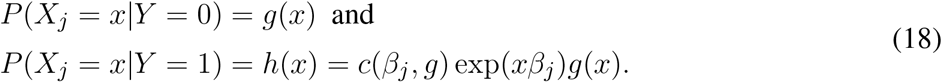

We assume that the distribution of *X* in controls, *g*(*x*), is Binomial(2, *p*), where *p* is the MAF of the SNV. Then the distribution of *X* in cases, *h*(*x*), is proportional to

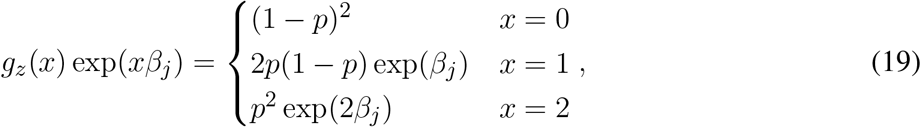

which has normalizing constant (1 − *p*)^2^ + 2*p*(1 − *p*) exp(*β*_*j*_) + *p*^2^ exp(2*β*_*j*_).

We simulated data in the presence of population stratification. We create population-disease and population-SNV associations as follows. To create population-disease association we introduced a population main effect on disease risk by taking population-stratum log-OR, *γ*, to be 1. To create population-SNV association we selected different SNV MAFs in different populations. Let *Z* denote a binary indicator of one of the two population strata. The respective frequencies in controls of the two populations are *f*_0_ and *f*_1_, respectively. Then the distribution of *Z* in controls is *P* (*Z* = *z*|*Y* = 0) = *f*_*z*_, and the distribution of *Z* in cases is *P* (*Z* = *z*|*Y* = 1) ∝ *f*_*z*_ exp(*zγ*) [31]. In our studies, we set *f*_0_ = *f*_1_ = 0.5. Now suppose that the MAF for a given SNV differs by sub-population, with *p*_*z*_ denoting the MAF in population *z*. Let *g*_*z*_(*x*) denote the distribution of *X*_*j*_ in controls of population *z*, i.e., *P* (*X*_*j*_ = *x*|*Z* = *z, Y* = 0) = *g*_*z*_(*x*) ∼ Binomial(2, *p*_*z*_). The joint distribution of *X* and *Z* in controls is then *P* (*X*_*j*_ = *x, Z* = *z*|*Y* = 0) = *f*_*z*_*g*_*z*_(*x*). If logit[*P* (*Y* = 1|*Z* = *z, X*_*j*_ = *x*)] = *α* + *zγ* + *xβ*_*j*_, the joint distribution of *X* and *Z* in cases is *P* (*X*_*j*_ = *x, Z* = *z*|*Y* = 1) ∝ *f*_*z*_*g*_*z*_(*x*) exp(*zγ* + *xβ*_*j*_) [15]. We then have

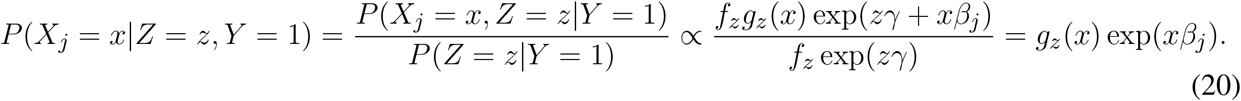

To summarize, we first assigned population status for each subject using

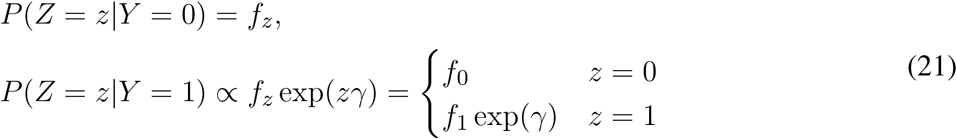

Then using (19), we simulated the genotype data of each SNV_*j*_, for *j* = 1, …, 1000, conditional on population status by sampling from

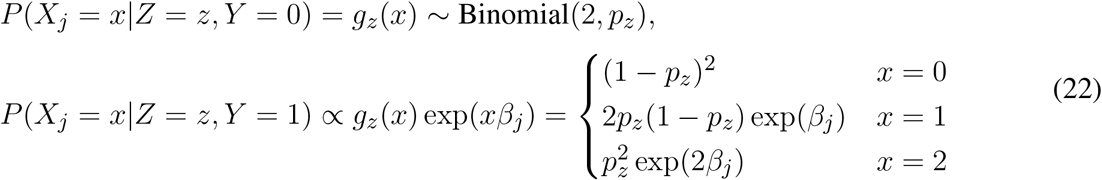

MAFs, *p*_*z*_, for different populations were obtained from 1000 Genomes Project [32]. Here we consider two populations: Caucasian (CEU) and Yoruba (YRI) subjects, and we sampled MAFs of SNVs from a 1 million base-pair region on Chromosome 6 (SNVs with MAF = 0 have been removed). Data from the 1000 Genomes Project was downloaded using the Data Slicer (https://www.internationalgenome.org/data-slicer/). The effect sizes of causal SNVs were assumed to be a decreasing function of MAF, which allows rare SNVs to have larger effect sizes and common SNVs to have smaller effect sizes. We set the magnitude of each 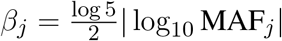 [21], where MAF_*j*_ is the pooled-MAFs (*p*_0_ + *p*_1_)*/*2 of the SNV_*j*_. We took into account the effects of mixed signs, multiplying *β*_*j*_ by -1 for some *j*, in which are 50% positive and 50% negative. This process gives the maximum OR = 6.44 (|*β*_*j*_| = 1.86) for SNVs with pooled-MAF = 0.0048 and the minimum OR = 1.40 (|*β*_*j*_| = 0.4) for SNVs with pooled-MAF = 0.38 (Supplementary Figure 2 B).

### 3.2 Results

We first evaluated the performance of the two different log-F methods described above. Over 100 simulation replicates, the mean estimates of *m* obtained by MCEM and LA are 4.77 (SD = 1.27) and 4.76 (SD = 1.18) respectively for *n* = 500, and are 3.88 (SD = 1.56) and 3.83 (SD = 1.33) respectively for *n* = 1000. The scatterplots (Figure 4) show good agreement between the two methods. Figure 5 compares the LA- and MCEM-based likelihood curves of *m* for the first 20 simulated data sets. These likelihoods were plotted with *m* of grid values from 1 to 10 on the x-axis, and each was smoothed by a smoothing spline. The likelihood curves are of similar shape, though shifted because the MCEM approach estimates the likelihood up to a constant (compare equations (13) and (14)). The compute time of LogF and FLR is given in Table 1. We see that LA is 160× and 300× faster than MCEM in elasped time for Step 1 when analyzing 1000 SNVs of sample size 500 and 1000, respectively. Although MCEM is computationally more expensive than LA, the accuracy of its approximation can be controlled by the number of Monte Carlo replicates, whereas the accuracy of LA cannot be controlled. We used *N* = 1000 Monte Carlo replicates in the MCEM throughout, which gives reasonably good accuracy. The agreement of the MCEM and LA approaches for smaller sample sizes validates the accuracy of LA. MCEM results are not available for the largest sample size of *n* = 1500, because our current implementation fails due to numerical underflow. As expected, once *m* is selected, LogF is computationally efficient as only a simple data augmentation approach is used in Step 2. Combining Step 1 (with LA) and Step 2, along with the preliminary scan, which is of the same order of computation time as Step 2, the combined computation time of the LogF approach is roughly half that of FLR.

**Table 1.** Computation time (elapsed time in seconds) of LogF and FLR when analyzing 1000 SNVs with sample size 500, 1000 and 1500 using 100 simulated data sets. No results are available for LogF-MCEM when *n* = 1500 due to numerical underflow.

| Method | Step | Min. | 1st Qu. | Elapsed time (s) |  | 3rd Qu. | Max. |
| --- | --- | --- | --- | --- | --- | --- | --- |
|  |  |  |  | Median | Mean |  |  |
| *n* = 500 |  |  |  |  |  |  |  |
| **LogF-MCEM** | **1** | 41 | 89 | 109 | 120 | 127 | 509 |
| **LogF-LA** | **1** | 0.35 | 0.49 | 0.54 | 0.74 | 0.58 | 13.22 |
| **LogF** | **2** | 3.53 | 3.66 | 3.69 | 3.78 | 3.92 | 4.31 |
| **FLR** | **NA** | 10.48 | 10.94 | 11.09 | 11.18 | 11.36 | 12.36 |
| *n* = 1000 |  |  |  |  |  |  |  |
| **LogF-MCEM** | **1** | 361 | 491 | 542 | 607 | 601 | 1274 |
| **LogF-LA** | **1** | 1.02 | 1.51 | 1.68 | 1.79 | 1.82 | 3.73 |
| **LogF** | **2** | 4.38 | 4.46 | 4.55 | 4.59 | 4.66 | 5.17 |
| **FLR** | **NA** | 17.04 | 17.60 | 17.79 | 17.85 | 18.00 | 19.48 |
| *n* = 1500 |  |  |  |  |  |  |  |
| **LogF-LA** | **1** | 2.42 | 2.66 | 2.75 | 2.78 | 2.89 | 3.73 |
| **LogF** | **2** | 5.21 | 5.30 | 5.37 | 5.39 | 5.39 | 5.94 |
| **FLR** | **NA** | 23.45 | 23.96 | 24.35 | 24.33 | 24.61 | 25.35 |

**Figure 4.**
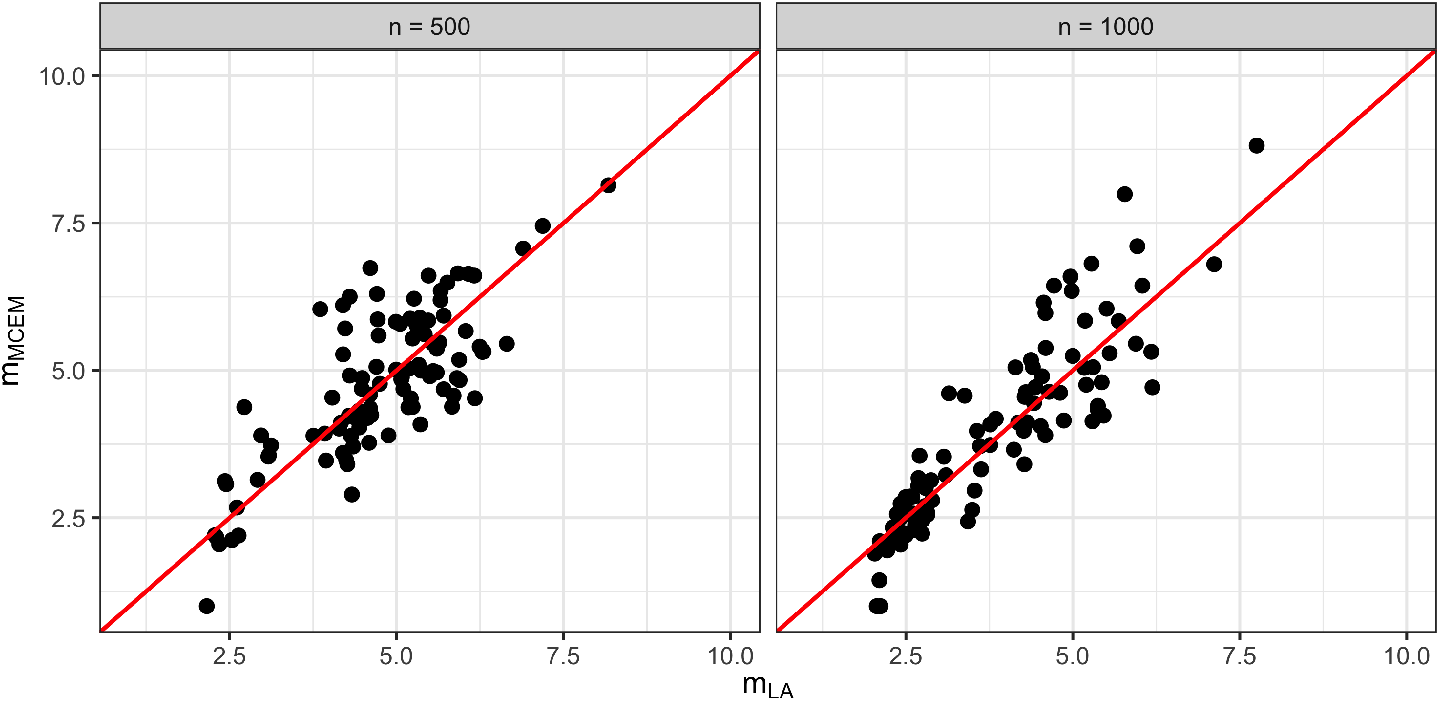
Scatterplot comparing the estimated values of *m* using the two methods over 100 simulation replicates. Values estimated by LA are on x-axis, and values estimated by MCEM are on y-axis. Red line is *y* = *x*.

**Figure 5.**
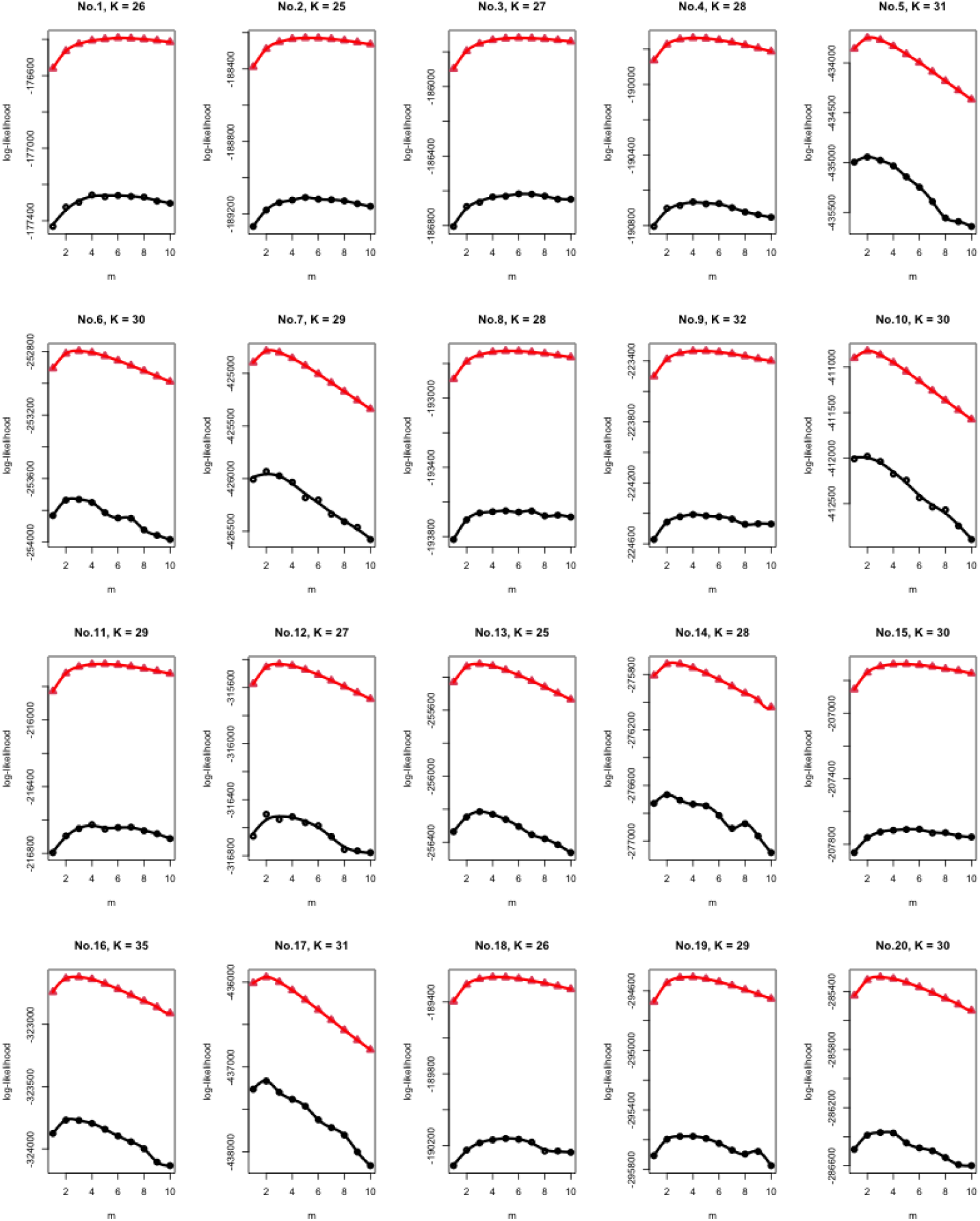
Comparison of profile log-likelihood curves obtained by the two different methods described in text, for the first 20 simulated data sets (*n* = 1000). In each case the likelihood curve was generated based on *K* SNVs selected in a preliminary genome-wide scan, and was smoothed by smoothing spline. The red line connecting triangles is based on LA, whereas the black line connecting dots corresponds to MCEM.

We further examined the accuracy of effect sizes from LogF-MCEM, LogF-LA, FLR and CP. All variants were binned based on the pooled-MAF in five bins: (0%, 1%), [1%, 5%), [5%, 10%), [10%, 25%) and [25%, 50%], and there were 51, 128, 213, 401, and 207 SNVs in each bin. The causal variants can be either deleterious or protective (i.e. *β*_*j*_ is either positive or negative), so we define the bias of effect size estimates as the signed bias, 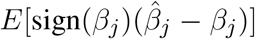; positive values indicate bias away from zero, while negative values indicate bias towards zero. We also evaluated the SD of effect size estimates as the standard deviation of 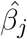 across 100 simulation replicates, and the mean squared error (MSE) as the sum of squared bias and squared SD. MAF-binned results are shown in Table 2 and Figure 6. In the Figure, results for the MLE obscure those for the other methods and are not shown. We find that for variants of MAF 1% or greater, all methods are comparable. However, for rare variants of MAF *<* 1%, the SD of LogF is much smaller than other methods. In addition, the signed bias of the LogF is more concentrated around zero compared with other methods, though this tendency about zero is counteracted by some extreme negative signed biases that suggest over-shrinkage in some cases. The MAFs of the three SNVs that lead to these extreme negative signed biases (Figure 6) are 0.0048, 0.0072, and 0.0074, respectively. We note that 0.0048 was the smallest MAF in our simulated datasets. Combining bias and SD results in a much smaller MSE for the LogF than other methods. Comparing the results under samples sizes of 500, 1000, and 1500, one can see that penalization makes less of an impact as the sample size increases.

**Table 2.**
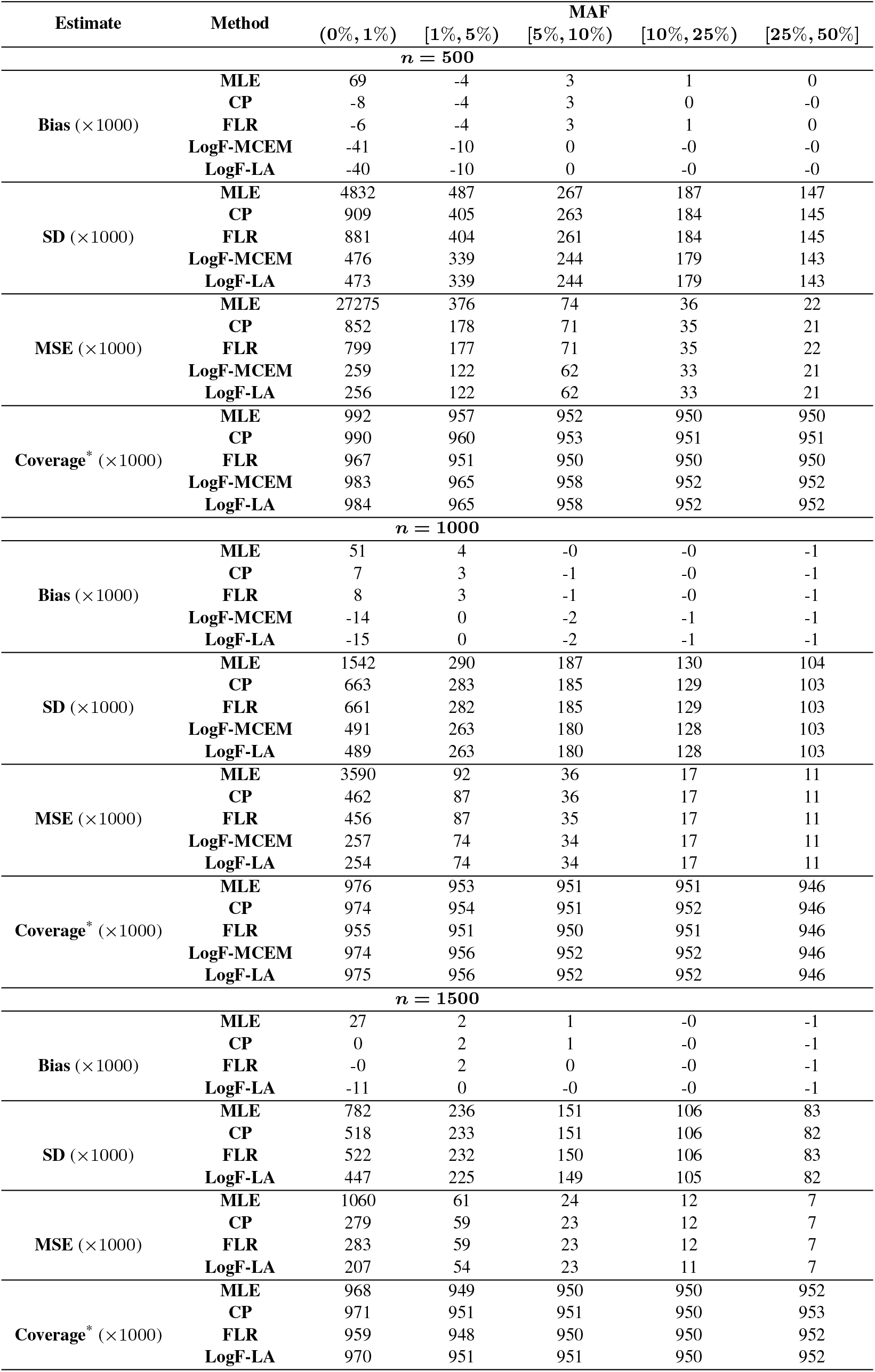
MAF binned averages of bias, SD, MSE and CI coverage probability of effect size estimates across 100 simulated data. * Coverage probability of two-sided nominal 95% confidence intervals for log-OR coefficient. Wald CIs were used for MLE and CP, whereas profile likelihood-based CIs were used for FLR, LogF-MCEM and LogF-LA. No results are available for LogF-MCEM when *n* = 1500 due to numerical underflow.

**Figure 6.**
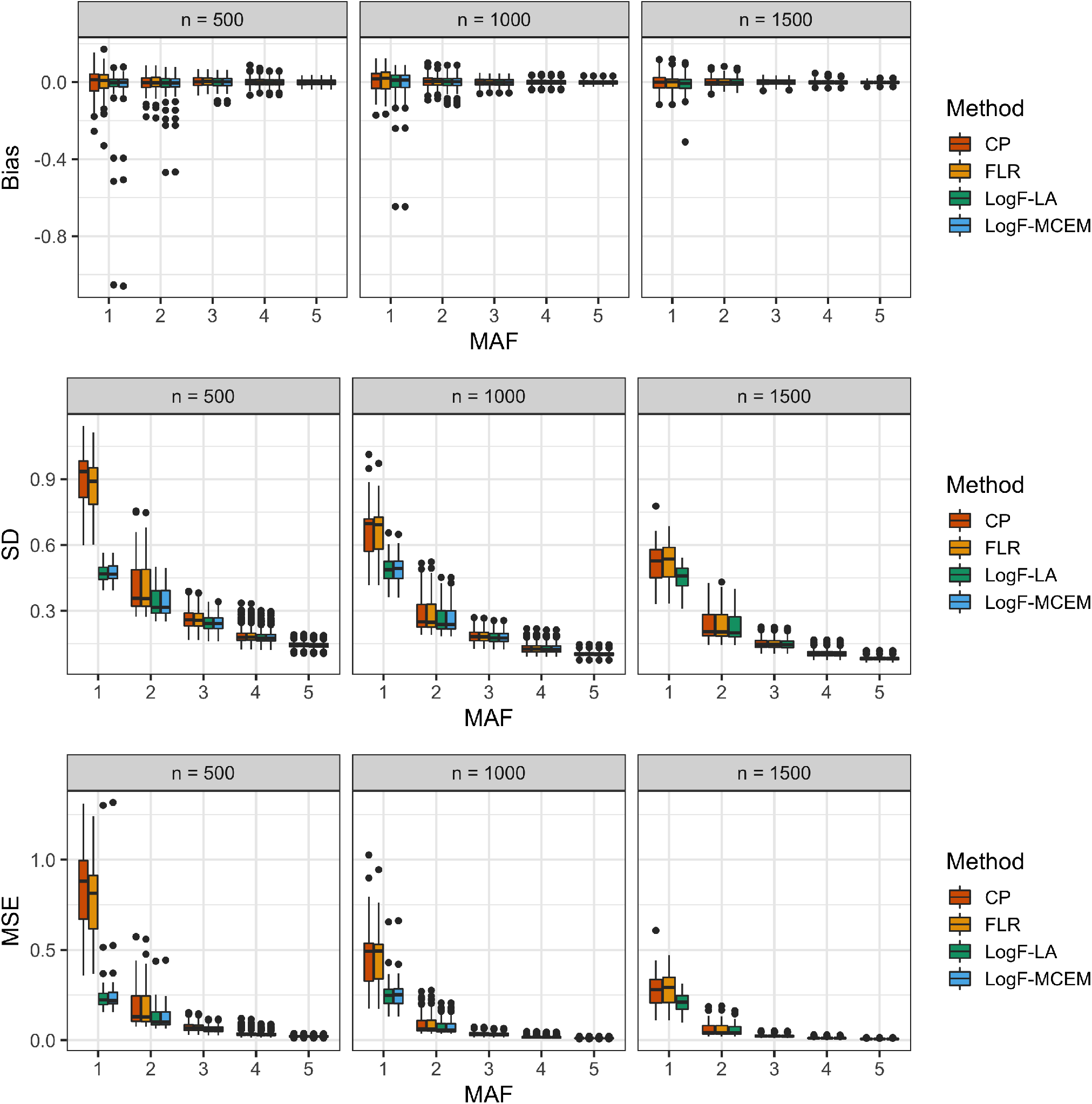
MAF binned boxplots of bias, SD and MSE of effect size estimates for LogF and other competing methods on simulated data. Each boxplot represents the distribution of the estimated quantity across 100 simulation replicates. MAF bins are: 1 = (0%, 1%), 2 = [1%, 5%), 3 = [5%, 10%), 4 = [10%, 25%) and 5 = [25%, 50%]. No results are available for LogF-MCEM when *n* = 1500 due to numerical underflow.

Through simulations, we also investigated the Type 1 error and power of the test of SNV effects from the different approaches (Figure 7). Although all the methods provide good control of Type 1 error, we found that LogF approaches result in a relatively smaller false positive rate. All the methods had similar power, with slightly less power from LogF approaches. We believe that the increased power of the FLR and Cauchy approaches can be partly attributed to their bias away from zero for rare variants.

**Figure 7.**
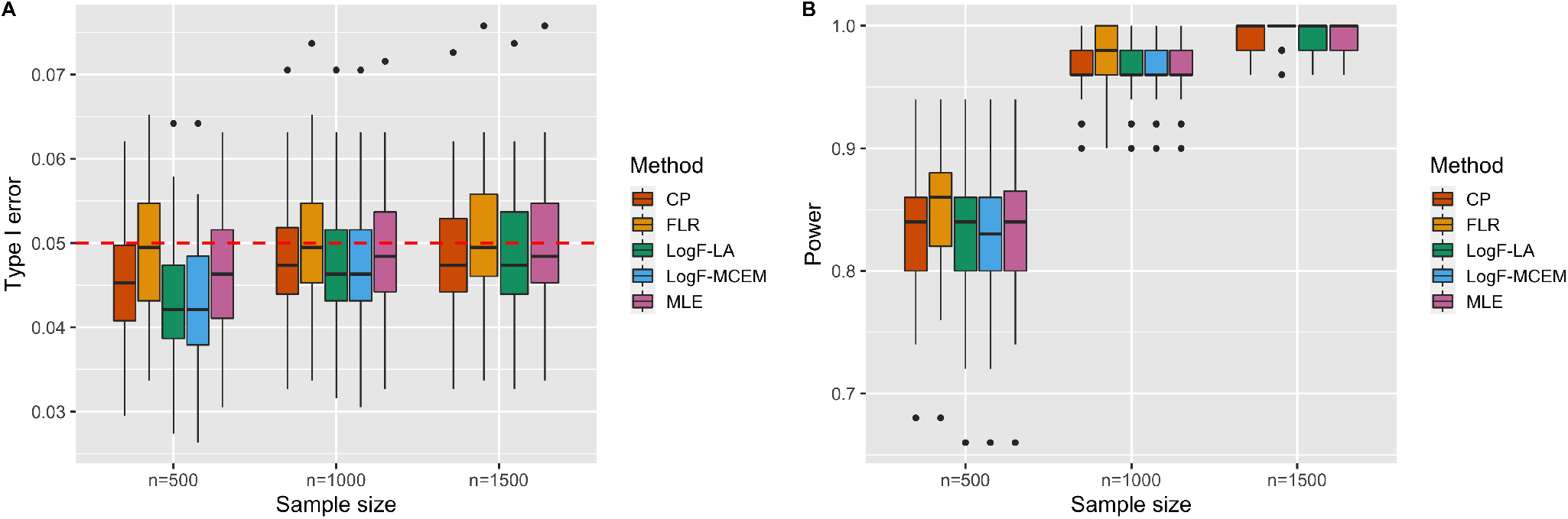
Type 1 error and power performance over simulated data sets. (A). Each boxplot represents the distribution of empirical type 1 error rates at nominal level 0.05 (red dashed horizontal line) across 100 simulation replicates computed at null SNVs. (B). Power computed at causal SNVs. No results are available for LogF-MCEM when *n* = 1500 due to numerical underflow.

## 4 DATA APPLICATION

The Super Seniors data from the Brooks-Wilson laboratory was collected to investigate the association between genetic heritability and healthy aging of humans. The ‘super seniors’ are defined as those who are 85 or older and have no history of being diagnosed with the following 5 types of diseases: cardiovascular disease, cancer, diabetes, major pulmonary disease or dementia. In this study, 1162 samples of 4,559,465 markers were genotyped using a custom Infinium Omni5Exome-4 v1.3 BeadChip (Illumina, San Diego, California, USA) at the McGill University/Genome Quebec Innovation Centre (Montreal, Quebec, Canada) [33]. The data underwent extensive quality control after genotyping, including re-clustering, removal of replicate and tri-allelic SNPs, and checking for sex discrepancies and relatedness. We also removed SNPs with MAF *<* 0.005, call rate *<* 97%, or Hardy-Weinberg equilibrium p-value *<* 1 × 10^−6^ among controls.

After a series of filtering steps, our final study includes 1044 self-reported Europeans, of which 427 are controls and 617 are cases (super seniors), and 2,678,703 autosomal SNPs.

A preliminary genome-wide scan identified 98 SNPs with p-values *<* 5 × 10^−5^. Of these, the 57 SNPs with no missing values were used to estimate the value of *m*. Our marginal likelihood approach for estimating *m* incorporates sex and the first 10 principal components as confounding variables. The *m* estimated by MCEM and LA are 7.01 and 6.89, respectively. To analyse 2,678,703 SNPs, the LogF approach (Step 2) takes 14 hours, which is 30× faster than FLR (437 hours). Manhattan plots (Figure 8) show very good agreement for the association detected between methods. Figure 9 shows the QQ-plot of p-values when applying MLE, LogF-LA (results of LogF-MCEM are close to LogF-LA, and are shown in Supplementary Figure 5-8) and FLR to the Super Seniors data. All methods are close to the dashed line of slope one, though the FLR p-values veer up slightly above the line at − log_10_ p-values near 5. Figure 10 compares the parameter estimates of the MLE, FLR and LogF. Other than cases where the MLE appears grossly inflated (e.g., 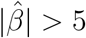), the estimates from the MLE and FLR are in surprisingly good agreement. The LogF estimates are shrunken more towards zero than those of FLR, and that the shrinkage is more pronounced for rare variants than for variants of frequency greater than 0.01. Figure 11 and Table 3 compare the p-values of the different approaches. The points below the dashed line of slope one in both panels of Figure 11 indicate that the FLR p-values are systematically lower than those of the MLE and LogF. This is also reflected in the confusion matrices of Table 3, which show that FLR flags more SNVs as significant at the 5 × 10^−5^ level than the other two methods. Taken together, these results suggest that the LogF approach may impose too much shrinkage on the SNV effect estimates.

**Table 3.**
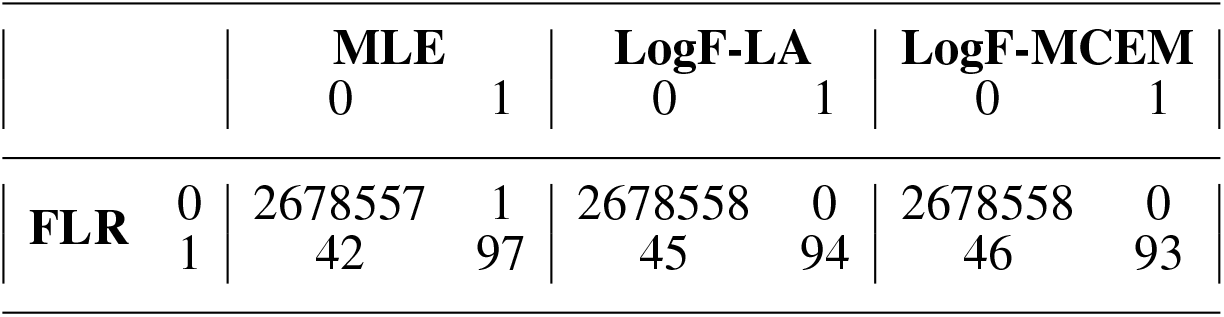
Confusion matrices comparing association results from different methods on Super Seniors data, where ’1’ indicating the number of SNPs below the genome-wide significant threshold of 5 × 10^−5^ and ’0’ otherwise.

**Figure 8.**
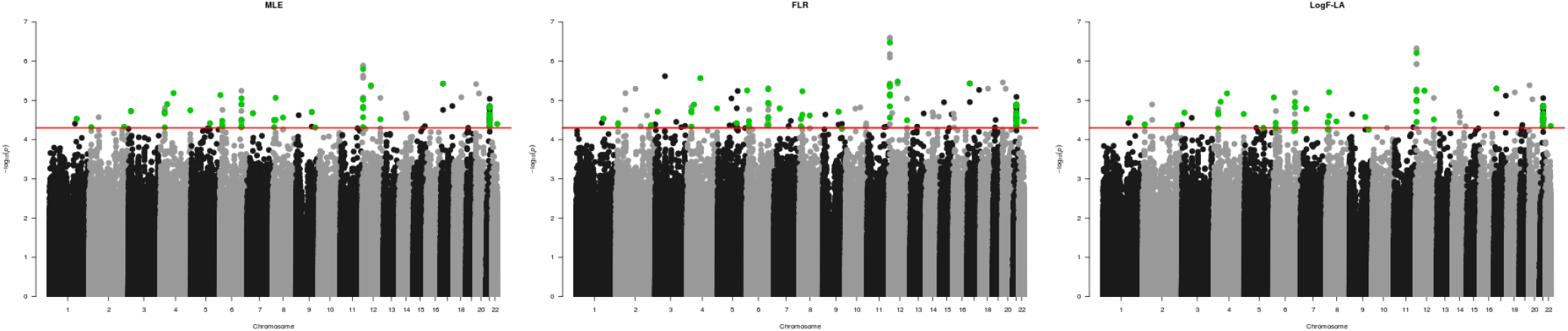
Manhattan plots comparing association results from different methods on Super Seniors data. The red horizontal line represents the liberal genome-wide significance threshold (*P* = 5 × 10^−5^) used to select SNPs in the preliminary scan. For LogF-LA, 57 SNPs (green points) below the threshold are used to estimate *m* in Step 1.

**Figure 9.**
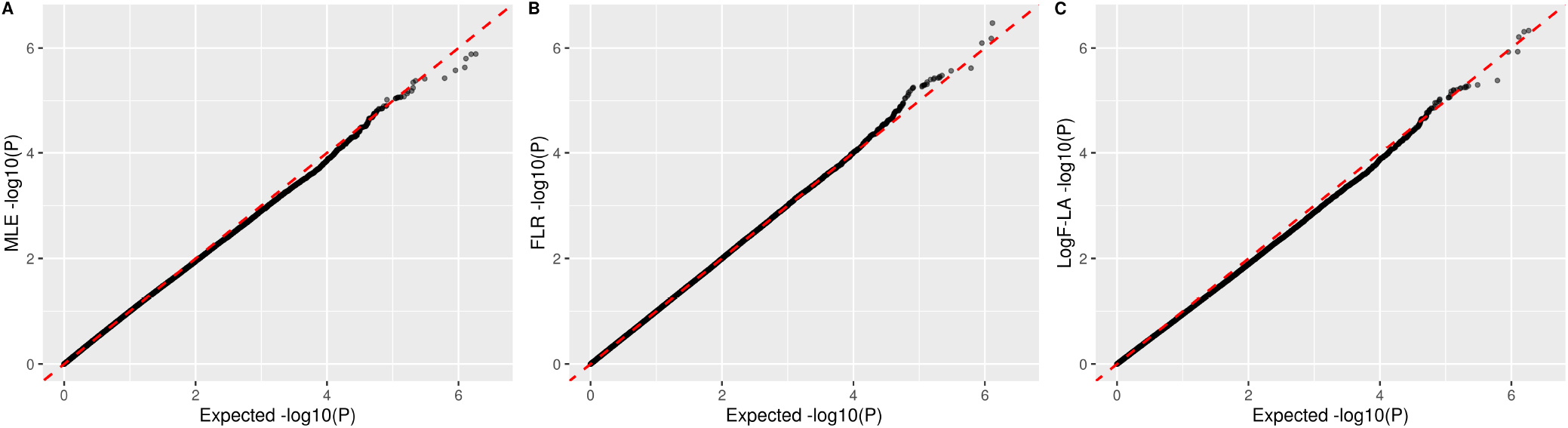
QQ-plot comparing p-values from different methods on Super Seniors data. The p-value for FLR and LogF-LA was obtained using the likelihood ratio test with a 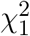 test statistic.

**Figure 10.**
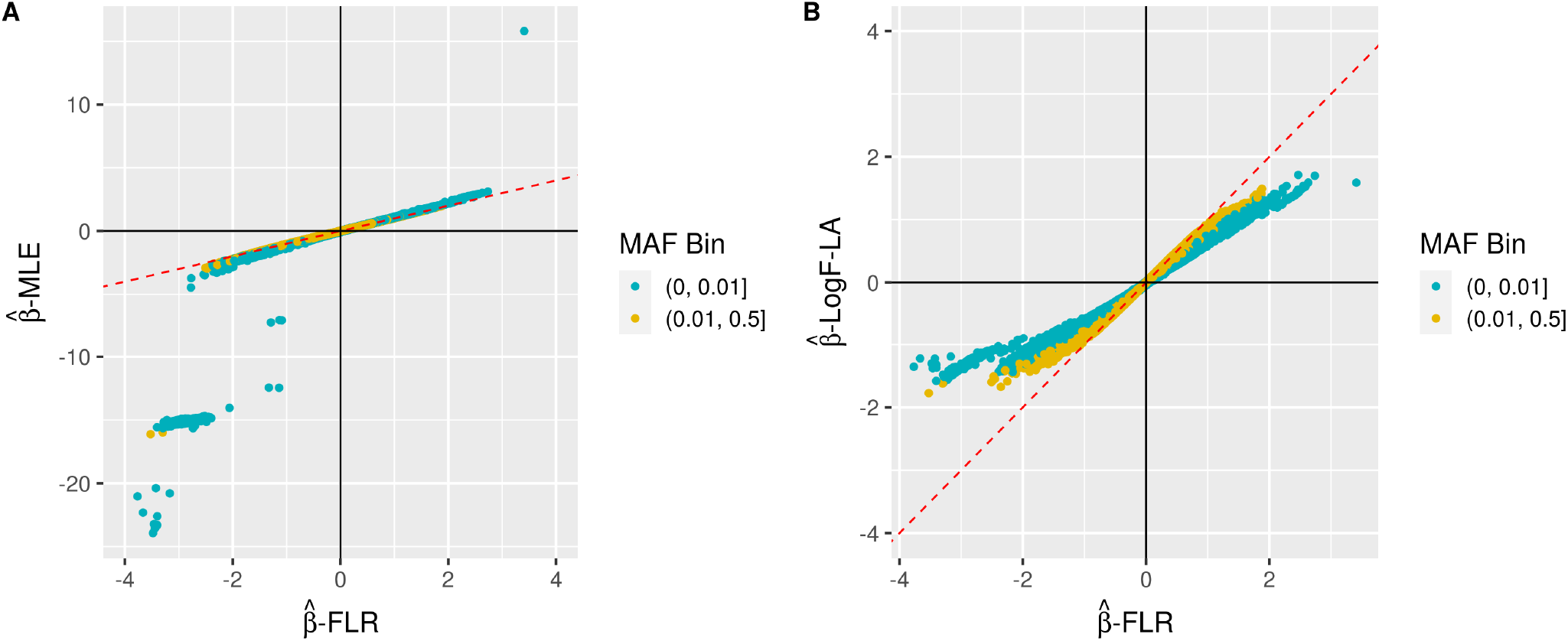
Scatterplots comparing effect size estimates from different methods for Super Seniors data. The plotting colors represents variant categories based on minor allele frequency (MAF) threshold of 1%.

**Figure 11.**
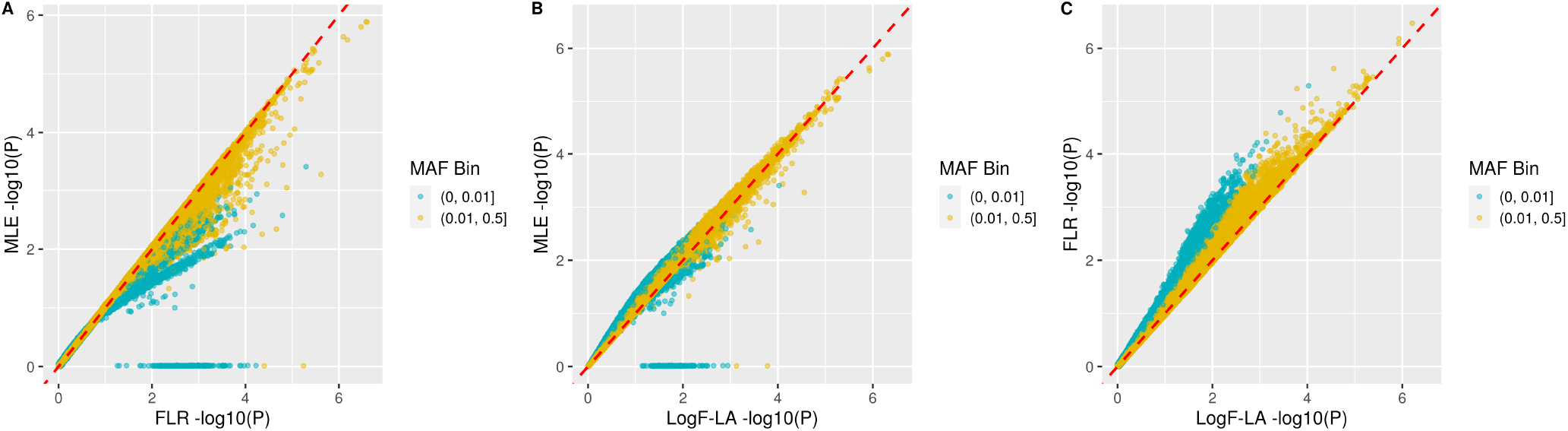
Scatterplots comparing p-values from different methods on Super Seniors data. The plotting colors represents variant categories based on minor allele frequency (MAF) threshold of 1%. For FLR and LogF-LA, the p-value for each variant was obtained by the likelihood ratio test with a 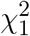 test statistic.

## 5 DISCUSSION AND CONCLUSION

We have proposed a method for single rare variant analysis with binary phenotypes by logistic regression penalized by log-*F* priors. Our approach consists of two steps. First, we select *K* markers that show evidence of association with the phenotype in a preliminary scan and use these to estimate *m*. The value of *m* is the maximizer of a composite of *K* marginal likelihoods obtained by integrating the random effect out of the joint distribution of the observed data and the random effect. Our maximization algorithm contains two approximate approaches: (1) a hybrid of an EM algorithm and brute-force maximization of Monte Carlo estimates of the marginal likelihood; and (2) a combination of a Laplace approximation and derivative-free optimization of the marginal likelihood. The two methods give similar results, with LA being faster for all sample sizes and more numerically stable for large sample sizes. Second, log-*F* penalties are conveniently implemented with standard logistic regression by translating the coefficient penalty into a pseudo-data record [12]. Our method requires extra computation time up front for the preliminary scan and selection of the shrinkage parameter *m*, but once selected, LogF approach (using LA in Step 1) is faster than Firth logistic regression (1). Our simulation studies suggest that the proposed LogF approach has slightly lower bias and substantially lower MSE than the other methods considered for variants of frequency less than 1%, and similar bias and MSE for variants of frequency greater than 1%. However, the power results of our simulation study and the analysis of the Super Seniors data suggest that our current implementation of log-*F* penalization has a tendency to over-shrink estimates of truly-associated SNVs. We discuss generalizations of the penalization approach that might correct such over-shrinkage in what follows.

Penalization can be generalized by allowing the prior distribution to depend on characteristics of the SNV, such as MAF or annotation information. A straightforward extension is to stratify selection of the shrinkage parameter by, e.g., MAF. That is, we might allow the prior distribution to be indexed by a variant-frequency-specific parameter instead of a common parameter for all variants. The idea could be as simple as multiplying the global shrinkage parameter *m* by a frequency-specific parameter *α*_*k*_; i.e., a variant in frequency bin *k* could have prior distribution log-*F* (*α*_*k*_*m, α*_*k*_*m*). We can choose the *α*_*k*_ values such that the distribution of common variants has a smaller variance and a larger variance for rare variants. In the context of heritability estimation [34] argue against stratified approaches and instead recommend modeling the variance of the SNV effects as proportional to [*f*_*i*_(1−*f*_*i*_)]^1−*α*^ for MAF *f*_*i*_ and a power *α*. Their analyses of real data suggested the value *α* = −0.25. This corresponds to standardizing each SNV covariate by dividing by [*f*_*i*_(1−*f*_*i*_)]^(1−*α*)*/*2^ before analyses. In the context of modelling quantitative traits [35], proposing a double-exponential prior on SNV effects and a log-linear model for the scale parameter of the double exponential distribution allows the scale to depend on SNV characteristics such as annotation information. We plan to investigate the properties of both standardization and modelling of the shrinkage parameter on data from the UK Biobank. We also plan to use the UK Biobank data to investigate how the shrinkage parameter depends on phenotype characteristics such as prevalence and heritability. Application of the LogF approach to data from the UK Biobank will also confirm that the methods scale to biobank-sized datasets.

It should be noted that in our simulations we used a simplified, binary confounding variable to represent population stratification. By contrast, the analysis of the Super Seniors data used an expanded set of confounding variables that included sex and 10 principal components. We have also mentioned adjustment for relatedness and population stratification by inclusion of an estimated polygenic effect as an “offset” in the model. Another extension of interest is to use log-*F* penalization for a SNV covariate of interest in a model that uses linear mixed models (LMMs) to correct for confounding due to population structure and genetic relatedness [36, 37]. LMMs can be viewed as regression with correlated errors, using a kinship matrix derived from anonymous SNVs to model correlations. It should be straightforward to extend this regression approach to include log-*F* penalization of the SNV of interest through data augmentation. Investigation of the properties of our approach in conjunction with LMMs is an area for future work.

In practice, identifying rare genetic causes of common diseases can improve diagnostic and treatment strategies for patients as well as provide insights into disease etiology. Recent studies have found that patients with low genetic risk scores (GRS) are more likely to carry rare pathogenic variants [38]. Although GRS are currently based on common variants, our method might be of use in extending GRS methods to include low-frequency or even rare variants of large effect sizes.

Our focus has been on single-SNV logistic regression, but log-*F* penalization generalizes to multiple-variant logistic regression. In general, we multiply the likelihood by as many log-*F* distributions as there are covariates whose coefficient we wish to penalize. This can also be implemented by a generalization of the data augmentation procedure described in Section 2.6 [12, 39]. Such an approach may be useful for performing the kinds of gene- or region-based tests that are commonly performed for rare variants, and investigation of its properties is ongoing.

## 6 STATEMENTS

A preprint version of this article is available on bioRxiv [40].

## 6.1 Acknowledgment

The authors have no acknowledgment to declare.

## 6.2 Statement of Ethics

The Super Seniors study was approved by the joint Clinical Research Ethics board of BC Cancer and The University of British Columbia. All participants provided written informed consent.

## 6.3 Conflict of Interest Statement

The authors have no conflicts of interest to declare.

## 6.4 Funding Sources

This work was supported, in part, by Discovery Grant RGPIN-05595 to Brad McNeney from the Natural Sciences and Engineering Research Council of Canada (NSERC). The Super Seniors study was established with a grant from the Canadian Institute of Health Research. Super Seniors genotype data generation and preparation were supported by grants from the Lotte and John Hecht Memorial Foundation and the Canadian Cancer Society.

## 6.5 Author Contributions

Ying Yu developed and implemented MCEM, generated simulated datasets, performed data application, and drafted the manuscript. Siyuan Chen developed and implemented LA. Brad McNeney developed the statistical methods and drafted the manuscript. Samantha J. Jones, Rawnak Hoque, Olga Vishnyakova, and Angela Brooks-Wilson prepared and QC’d the Super Seniors data. All authors revised the manuscript and approved the final version.

## 6.6 Data Availability Statement

Datasets simulated to evaluate the properties of the proposed method are available on request from the corresponding author. R code to implement the methods is available from https://github.com/SFUStatgen/logistlogF.

## FIGURE LEGENDS

- Supplementary Fig. 1. ***Y***_(*Nn*×1)_ is a vector containing *N* replicates of ***y*** and ***X***_(*Nn*×1)_ is a vector containing *N* replicates of ***x. W*** stands for the weights for each Monte Carlo replicate such that 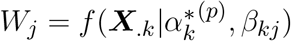 and the offset term 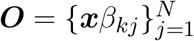.
- Supplementary Fig. 2. A. Histogram of 1000 SNV-effect-sizes used for data simulation, in which are 5 casual SNVs and 950 null SNVs. B. Histogram of effect sizes of causal SNPs, where 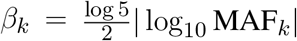.
- Supplementary Fig. 3. Effect sizes of 1000 SNVs generated used for data simulation by minor allele frequency. Red dots indicate casual SNVs and blue dots indicate non-casual SNVs.
- Supplementary Fig. 4. Manhattan plots showing association results from LogF-MCEM on Super Seniors data. The red horizontal line represents the liberal genome-wide significance threshold (*P* = 5 × 10^−5^) used to select SNPs in the preliminary scan. 57 SNPs (green points) below the threshold are used to estimate *m* in Step 1.
- Supplementary Fig. 5. QQ-plot showing p-values from LogF-MCEM on Super Seniors data. The p-value was obtained using the likelihood ratio test with a 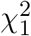 test statistic.
- Supplementary Fig. 6. Scatterplots showing effect size estimates from LogF-MCEM for Super Seniors data. The plotting colors represents variant categories based on minor allele frequency (MAF) threshold of 1%.
- Supplementary Fig. 7. Scatterplots comparing p-values from different methods on Super Seniors data. The plotting colors represents variant categories based on minor allele frequency (MAF) threshold of 1%. For FLR and LogF-MCEM, the p-value for each variant was obtained by the likelihood ratio test with a 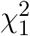 test statistic.

## Notes

https://github.com/SFUStatgen/PenalizedLR

### Competing Interest Statement

The authors have declared no competing interest.

### Summary of Updates

Manuscript updated

